# Metaplastic sleep regulation in *Drosophila* determined by microscale circadian neural dynamics

**DOI:** 10.64898/2026.03.21.713346

**Authors:** Anelise N. Hutson, Alexa N. Zarjetskiy, Yutian J. Zhang, Natalia Pokaleva, Elizabeth M. Paul, Yizhou Xie, Bryan Chong, Victor M. Sanchez Franco, Lauren H. Zukowski, Eileen E. Faulk, Joel A. Walker, Ayanna M. Brown, Dieu Linh Nguyen, Faith S. Ferry, Eleanora M. Snyder, Masashi Tabuchi

## Abstract

The biophysical mechanisms by which circadian clock neurons integrate temporal coding signals to regulate sleep remain elusive. Here, using *Drosophila*, we identify Rabphilin (Rph) in DN1p clock neurons as a key stabilizer of the metaplasticity setpoint governing circadian regulation of sleep. Rph protein levels are elevated at night relative to daytime and modulate stochastic process of DN1p membrane potential dynamics linked to variability in synaptic activity at connections between DN1p neurons and their downstream postsynaptic partners. We find that Rph acts as a bidirectional regulator of synaptic plasticity thresholds. Under dim nocturnal light stimulation, Rph knockdown permits synaptic potentiation, whereas synthesized Rph introduction induces synaptic depression. In contrast, under optogenetic manipulation mimicking daytime spiking in DN1p neurons, these effects are reversed. We further show that spike-timing–dependent plasticity emerges when postsynaptic spiking is engaged, with nocturnal dim light conditions determining the direction of plasticity. Together, these findings establish a mechanistic link between microscale circadian neural dynamics and hierarchical metaplastic regulation, demonstrating how circadian regulation of sleep dynamically balances stability and adaptive flexibility through circadian setpoints and environmental nocturnal light interactions.

**Significance Statement:** We show that circadian metaplasticity regulates sleep through membrane potential dynamics. Circadian clock neurons implement flexible metaplasticity, whereby the direction can be determined by internal circadian setpoints and interactions with nocturnal environmental light. This mechanism engages spike-timing–dependent plasticity to determine plasticity polarity. Our findings identify membrane potential dynamics as a computational substrate for physiological state control, linking molecular mechanisms to circuit-level circadian regulation of sleep. Together, they reframe sleep regulation as an active metaplastic process that hierarchically integrates microscale circadian neural dynamics to optimize circuit function.

## Introduction

A central challenge in neuroscience is understanding how neural circuits transform molecular signals into coherent behavioral outputs (1-5). Although the molecular architecture of circadian clocks has been extensively characterized, the computational principles that convert 24-hour molecular oscillations into precisely timed behaviors remain unresolved. Circadian rhythms are fundamental biological processes that coordinate physiological and behavioral functions across organisms, ensuring optimal adaptation to the 24-hour environmental cycle (6-9). While this framework accounts for rhythmicity, it fails to explain two fundamental features of circadian behavior: the coexistence of temporal precision with environmental flexibility, and the capacity of identical molecular oscillations to produce context-dependent physiological and behavioral outputs. These limitations suggest the existence of an intermediate computational layer that has not yet been identified. Circadian regulation extends to multiple neural processes, with the suprachiasmatic nucleus in mammals and analogous structures in other organisms serving as master pacemakers that orchestrate physiologically important behaviors such as sleep regulation (10-12). In *Drosophila*, the electrical activity in DN1ps circadian clock neurons (13-16), play crucial roles in regulating sleep (17-19) by modulating electrical and synaptic activities in response to environmental cues (13, 20-23). However, despite significant advances in identifying the molecular identifications, including core clock genes that drive rhythms via transcriptional–translational negative feedback (24-27), as well as their output molecules (28, 29), the biophysical and computational principles by which molecular clocks generate neural codes remain largely understood (30, 31). This knowledge gap is particularly pronounced regarding the role of membrane potential dynamics and their stability in shaping synaptic plasticity. Identifying and characterizing these dynamics poses significant methodological challenges, requiring demanding intracellular electrophysiological recordings and computational signal processing to reveal subtle, hidden microscale temporal structures that may be emerged by environmental and/or internal noise (32-34). A framework to explore how synaptic plasticity can be utilized for sleep regulation (35), but how molecular clock dynamics are implemented in electrophysiological states to regulate sleep remains unclear. We conducted a large-scale RNAi screening to delineate the molecular basis of synaptic plasticity in DN1ps (21), we identified Rabphilin (Rph). Rph is a synaptic vesicle-associated protein that serves as a Rab3 effector and is implicated in synaptic regulation (36-40). We hypothesized that Rph controls synaptic plasticity by regulating microscale dynamical instabilities (41), thereby forming neural codes at DN1-PI synapses that govern circadian sleep regulation in *Drosophila*. Through genetic and electrophysiological analyses, we demonstrate that Rph governs metaplasticity by modulating the impact of nocturnal light exposure, which otherwise destabilizes neuronal dynamics and drives synaptic potentiation. This metaplastic mechanism aligns with the synaptic homeostasis hypothesis (35, 42), whereby sleep allows for global synaptic downscaling after wake-induced potentiation, thereby restoring learning capacity and preventing synaptic saturation, providing a conceptual advance in understanding how microscale membrane potential fluctuations profoundly influence the macroscale dynamics of circadian neurons and sleep-wake regulation.

## Results

### Rabphilin in DN1p clock neurons mediates sleep quality

To identify molecular regulators of sleep quality, we conducted a large-scale RNAi screen targeting 847 genes expressed in DN1p circadian clock neurons in *Drosophila*. This approach identified Rph as one of the strongest modulators of sleep architecture. Rph is a synaptic vesicle–associated protein that interacts with Rab3A and regulates calcium-dependent exocytosis through its C2 domains (43, 44); however, its role in circadian clock neurons has not been explored. DN1p-specific RNAi knockdown of Rph based on *R18H11-GAL4>UAS-Rph RNAi* flies caused exhibited altered sleep patterns compared to controls (**Fig. 1A-D and Fig. S1**), including increased brief awakening (**Fig. 1D**) suggesting fragmented sleep architecture. In addition to increased brief awakening, sleep bout duration was markedly reduced in flies expressing Rph RNAi in DN1p neurons (**Fig. S1B**), and this reduction in sleep bout duration was accompanied by increased bout number (**Fig. S1C**) and increased active time (**Fig. 1B**) with changes in activity intensity per minute during wakefulness (**Fig. 1C**). To examine whether Rph expression level has circadian cycling, we next performed western blotting. Western blotting analysis revealed that Rph protein levels in *Drosophila* heads exhibited robust circadian oscillation, with significantly higher expression during nighttime at ZT 18-20, compared to daytime at ZT 6-8 (**Fig. 1E-F**). Alpha-tubulin levels remained constant across time points, serving as an internal control (**Fig. 1G**). We also performed qPCR to quantify Rph mRNA in DN1p neurons, and found there is no significant difference between ZT6-8 and ZT18-20 (**Fig. 1H**), indicating circadian difference observed in western blotting could be based on post-transcriptional regulation of Rph. We also performed immunostaining by using anti-Rph antibodies, and fond that Rph is certainly expressed in DN1p neurons (**Fig. S1E**).

**Fig. 1.**
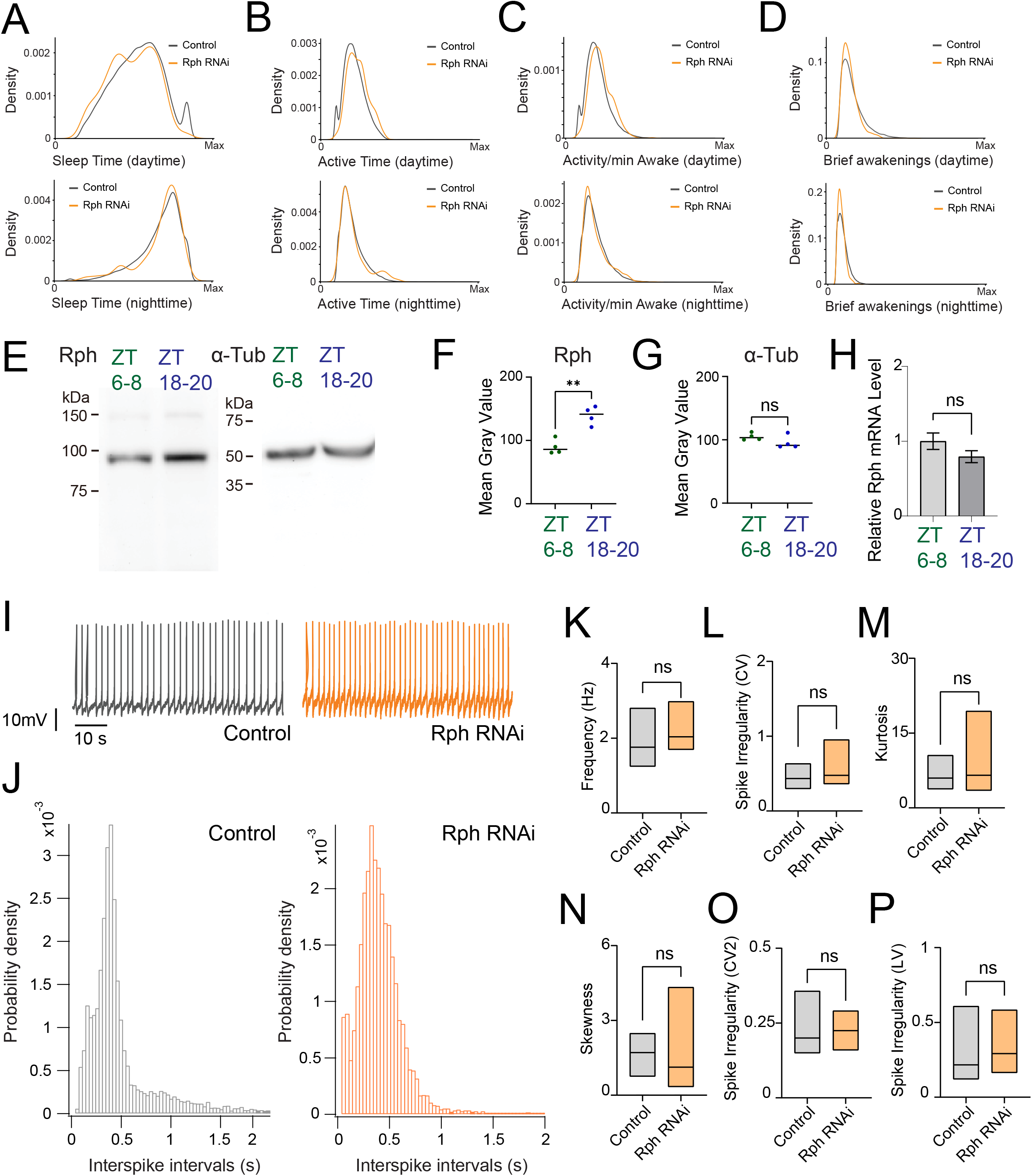
Rph regulates sleep, head Rph abundance and spontaneous activity in DN1p neurons. (**A–D**) Sleep phenotypes of *R18H11-GAL4>ctrl RNAi* flies (gray, N = 64) and *R18H11-GAL4>UAS-Rph RNAi #1* flies (orange, N = 128). (**A**) Total sleep during the day (ZT0–12) and night (ZT12–24). (**B**) Total time spent active during the day and night. (**C**) Activity per minute during wakefulness. (**D**) Number of brief awakenings during the day and night. (**E–H**) Western blot analyses of Rabphilin (Rph) in fly heads. (**E**) Representative immunoblots for Rph and α-tubulin from heads collected during the day (ZT6–8; green; N = 4) or night (ZT18– 20; blue; N = 4). (**F**) Quantification of Rph Western blot signals. (**G**) Quantification of α-tubulin Western blot signals. (**H**) qPCR showing relative Rph mRNA levels during ZT6-8 (N=4) or ZT18-20 (N=4). (**I–P**) Spontaneous firing properties of DN1p neurons at ZT18-20 in control flies and flies expressing Rph RNAi in DN1p neurons. (**I**) Representative membrane voltage traces of spontaneous activity of DN1p neurons at ZT18- 20. (**J**) Probability density distributions of interspike intervals. (**K**) Mean firing rate. (**L**) Coefficient of variation (CV) of interspike intervals. (**M**) Kurtosis of spontaneous firing distributions. (**N**) Skewness of spontaneous firing distributions. (**O**) Local coefficient of variation (CV2). (**P**) Local variation (LV). Sample sizes: control, N = 32 flies; Rph RNAi, N = 32 flies. The statistics used were an unpaired t-test with *p < 0.05, **p < 0.01, ***p < 0.001, ****p < 0.0001, and ns indicated non-significant.

### Rph selectively regulates subthreshold membrane dynamics without altering spike output in DN1p neurons

To define the cellular mechanism by which Rph contributes to circadian sleep regulation, we performed sharp-electrode intracellular recordings from DN1p neurons and quantified their spontaneous firing activity (**Fig. 1I**). We found that RNAi knockdown of Rph in DN1p neurons did not alter mean firing rate (**Fig. 1K**), conventional measures of spike-train variability, including the coefficient of variation, skewness and kurtosis of interspike intervals (**Fig. 1L–N**), or local irregularity metrics based on adjacent interspike-interval relationships (**Fig. 1O,P**). Current-injection protocols likewise revealed no detectable genotype-dependent differences in evoked firing (**Fig. S2**). Thus, Rph is not required for basal spike output or gross excitability in DN1p neurons. Because spike-based measures were preserved, we next asked whether Rph instead regulates subthreshold membrane behavior. To capture hidden temporal structure in membrane-voltage fluctuations, we modeled the recordings using an Ornstein–Uhlenbeck (OU) process. This analysis revealed that Rph RNAi knockdown increased the standard deviation of spontaneous membrane fluctuations and altered their decay dynamics, including both decay time and decay slope (**Fig. 2A–C**). Consistent with this, the OU noise amplitude (σ) was significantly elevated in Rph-deficient neurons (**Fig. 2D**), whereas the time constant (τ) showed a more modest shift (**Fig. 2E,F**), indicating that Rph primarily constrains the magnitude of spontaneous voltage fluctuations while also shaping their relaxation kinetics. Continuous wavelet transform analysis further showed a reorganization of frequency-dependent variability (**Fig. 2G,H**), and Augmented Dickey–Fuller testing identified genotype-dependent disruptions in stationarity at frequencies above 59 Hz (**Fig. 2I**). At the behavioral level, we found that these electrophysiological changes in stochastic process of DN1p membrane potential dynamics were accompanied by increased nighttime movement activity (**Fig. 2J**), resembling the phenotype observed under constant-light conditions. Together, these data identify Rph as a selective regulator of stochastic process of subthreshold membrane potential dynamics, rather than spike generation, in DN1p neurons.

**Fig. 2.**
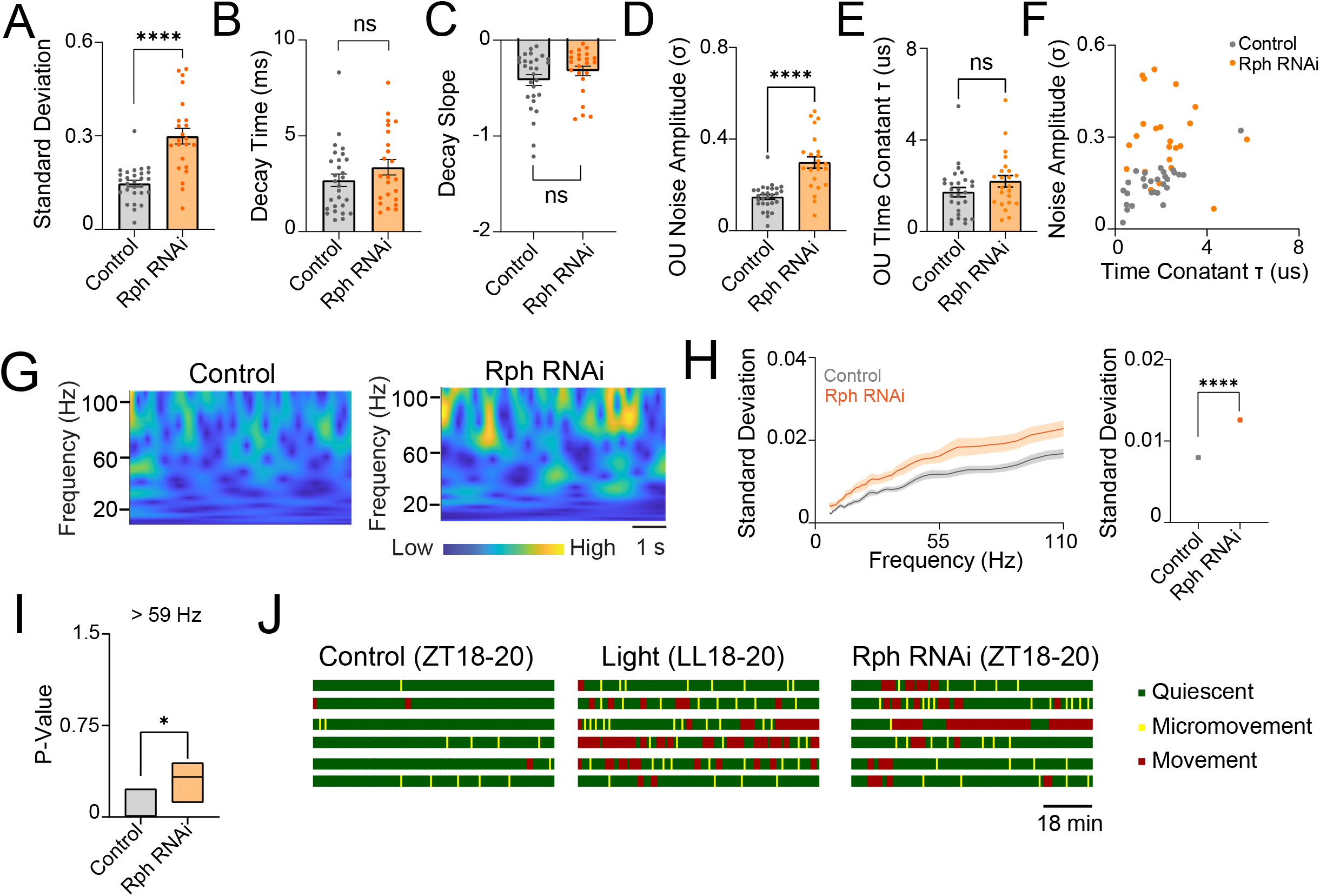
Ornstein–Uhlenbeck modeling of DN1p membrane voltage dynamics and nighttime movement. (**A–C**) Ornstein–Uhlenbeck (OU) fits derived from electrophysiological recordings of membrane potential of spontaneous activity of DN1p neurons at ZT18-20 in control flies (gray, N = 28) and flies expressing Rph RNAi in DN1p neurons (orange, N = 23). (**A**) Variability of the time series of the OU process. (**B**) Decay time constant. (**C**) Decay slope. (**D–I**) Statistical and spectral parameters of the OU model. (**D**) OU noise amplitude (σ), defined as the variability of membrane-voltage fluctuations. (**E**) OU time constant (τ), defined as the exponential decay constant of the autocorrelation function. (**F**) OU noise amplitude plotted against the corresponding OU time constant for each recording. (**G**) Continuous wavelet transform (CWT) of OU-model time series derived from membrane-voltage recordings. (**H**) Standard deviation of the CWT as a function of frequency and averaged across all frequencies. (**I**) Stationarity of OU-modeled time series assessed by Augmented Dickey–Fuller (ADF) tests; P values for frequencies greater than 59 Hz are shown for control flies and flies expressing Rph RNAi in DN1p neurons. (**J**) Single-fly video tracking of movement at ZT18-20 in control flies, flies maintained in constant light at LL18-20, and flies expressing Rph RNAi in DN1p neurons. The statistics used were an unpaired t-test with *p < 0.05, **p < 0.01, ***p < 0.001, ****p < 0.0001, and ns indicated non-significant.

### Rph-dependent changes in DN1p are transmitted to downstream PI neurons as altered synaptic variability

To determine whether altered DN1p membrane dynamics propagate to downstream sleep circuitry, we recorded spontaneous postsynaptic potentials (PSPs) from neurons in the pars intercerebralis (PI), a key target of DN1p output. Relative to controls at ZT18-20, both constant-light exposure at LL18-20 and Rph RNAi knockdown in DN1p neurons at ZT18-20 (**Fig. S3A**) altered the distribution of spontaneous PSPs in PI neurons, changing PSP amplitude variability, PSP slope and inter-event interval structure (**Fig. 3A–D**). Similar changes were also observed in the mPSP analysis. (**Fig. S3B-G**). Thus, perturbing either environmental circadian input or Rph- dependent molecular state in DN1p neurons produced convergent effects on the statistics of postsynaptic activity. We next asked whether these changes reflected a reorganization of synaptic-state transitions rather than a simple shift in mean PSP size. K-means clustering separated spontaneous events into larger PSPs and miniature PSPs (**Fig. S3B**), and discrete-time Markov chain analysis (45) revealed that the transition probabilities between these states were altered by both constant light and Rph knockdown. Transition probability analysis revealed an elevated probability of PSP→mPSP state transitions and a corresponding reduction in PSP→PSP transitions (**Fig. 3E**), suggesting enhanced temporal correlation among spontaneous release events. This result indicates that Rph influences the temporal organization of synaptic drive to PI neurons. Acute addition of exogenous Rph shifted stimulus-evoked changes in PSP slope toward the control state (**Fig. 3F**), supporting a causal role for Rph in determining synaptic response gain. Because PI neurons are bilaterally organized, we also tested whether Rph affects coordination across hemispheres. Dual-electrode recordings revealed prominent near-simultaneous PSP events in control flies (**Fig. 3G**). Both constant-light exposure and Rph knockdown reduced correlation coefficient and probability of cross-synaptic synchrony between the left and right PI (**Fig. 3H,I and Fig. S4A**), indicating that Rph contributes not only to synaptic variability but also to the bilateral coherence of circuit output.

**Fig. 3.**
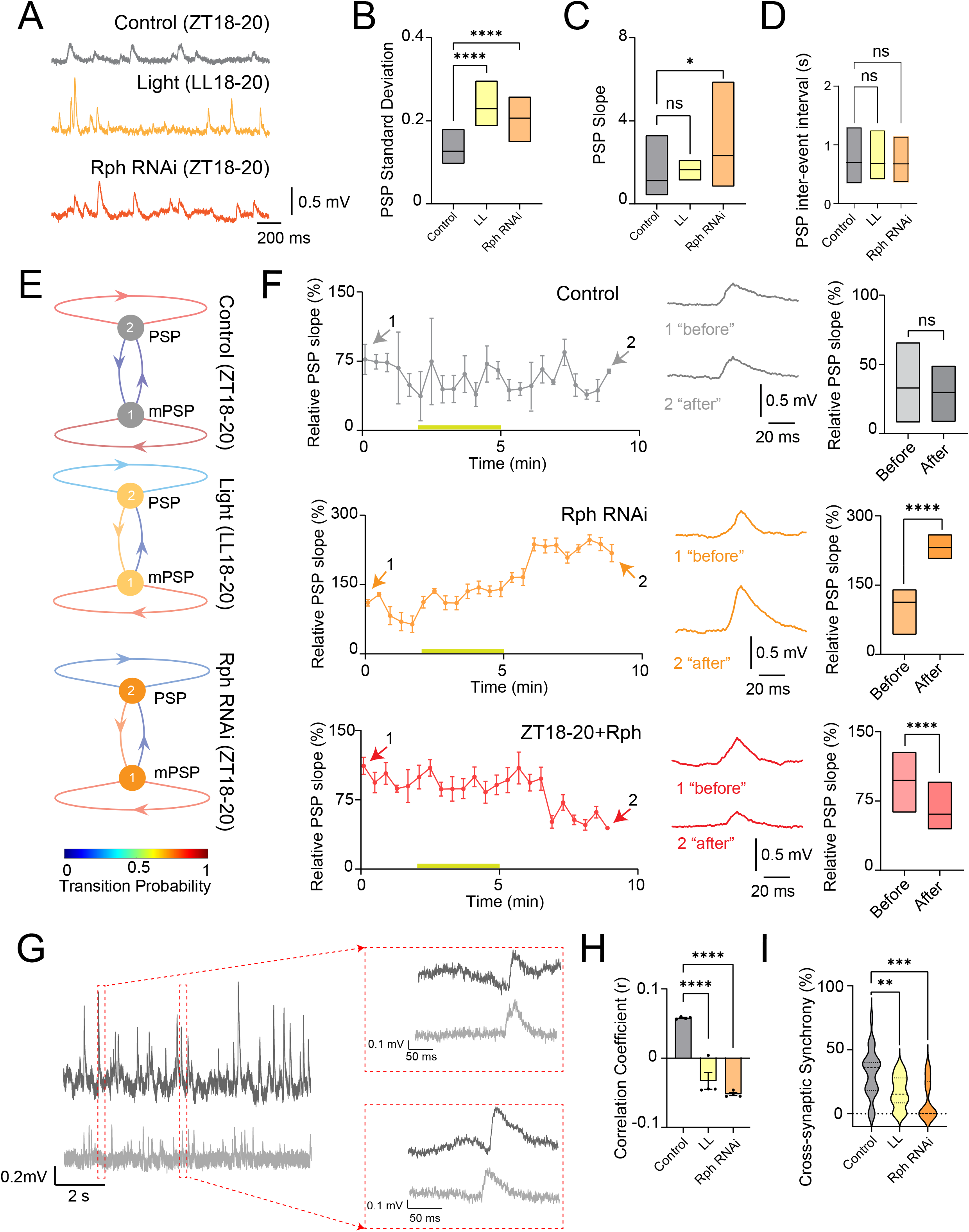
Rph in DN1p neurons regulates spontaneous synaptic variability and bilateral synchrony in downstream pars intercerebralis neurons. (**A–F**) Spontaneous postsynaptic potentials (PSPs) recorded from pars intercerebralis (PI) neurons. (**A**) Representative membrane voltage traces of spontaneous PSPs in PI neurons. (**B**)Variability of PSP amplitudes in PI neurons from control flies at ZT18-20 (gray, N = 8), flies maintained in constant light at LL18-20 (yellow, N = 8), and flies expressing Rph RNAi in DN1p neurons at ZT18-20 (orange, N = 8). (**C**) PSP slope. (**D**) Inter-event interval of PSPs. (**E**) State-transition analysis of PSP amplitudes using discrete-time Markov chains. PSP events were classified by *k*-means clustering into two states, PSPs and miniature PSPs (mPSPs), and the corresponding transition matrix was used to estimate class-transition probabilities. (**F**) Time course of relative PSP slope before, during and after stimulation in control flies at ZT18-20 (gray, N = 14), flies expressing Rph RNAi in DN1p neurons at ZT18-20 (orange, N = 15), and after synthesized Rph addition at ZT18-20 (red, N = 14). Quantification of the mean relative slope before and after stimulation is shown at right. The statistics used were an unpaired t-test with *p < 0.05, **p < 0.01, ***p < 0.001, ****p < 0.0001, and ns indicated non-significant. (**G–I**) Dual-electrode recordings assessing cross-synaptic coupling between the left and right PI. (**G**) Representative examples of near-synchronous PSPs in control flies at ZT18-20. (**H**) Correlation coefficient between hemispheres in control flies at ZT18-20 (N=4), flies maintained in constant light at LL18-20 (N=4) and flies expressing Rph RNAi in DN1p neurons at ZT18-20 (N=4). (**I**) Percentage of cross-synaptic synchrony between hemispheres in control flies at ZT18-20 (N=4), flies maintained in constant light at LL18-20 (N=4) and flies expressing Rph RNAi in DN1p neurons at ZT18-20 (N=4). **p < 0.01, ***p < 0.001, ****p < 0.0001. ns: non-significance based on one-way ANOVA followed by post-hoc Tukey tests.

### Rph sets the threshold for homosynaptic plasticity in PI neurons

The changes in spontaneous PSP statistics suggested that Rph might regulate the metaplastic state of the DN1p→PI circuit. We therefore investigated stimulus-evoked synaptic plasticity using optogenetic manipulation modeled on DN1p “daytime” spiking patterns, previously demonstrated to induce synaptic potentiation in PI neurons at ZT18–20 (21). In control flies, as shown previously (21), using optogenetic manipulation induced robust synaptic potentiation, evident as a sustained increase in relative PSP slope after induction (**Fig. 4A**). By contrast, the same using optogenetic manipulation protocol in flies exposed to constant light produced synaptic depression (**Fig. 4B**) at LL18-20, indicating that environmental light disruption of circadian state shifts the sign of synaptic modification. Strikingly, selective RNAi knockdown of Rph in DN1p neurons phenocopied the constant-light condition: the same stimulation protocol now induced synaptic depression rather than synaptic potentiation (**Fig. 4C**). Conversely, addition of exogenous Rph to preparations from constant-light flies shifted the response back toward the control state (**Fig. 4D**). These findings show that Rph does not simply increase or decrease synaptic strength; rather, it acts as a threshold-setting factor that determines whether identical input drives potentiation or depression.

**Fig. 4.**
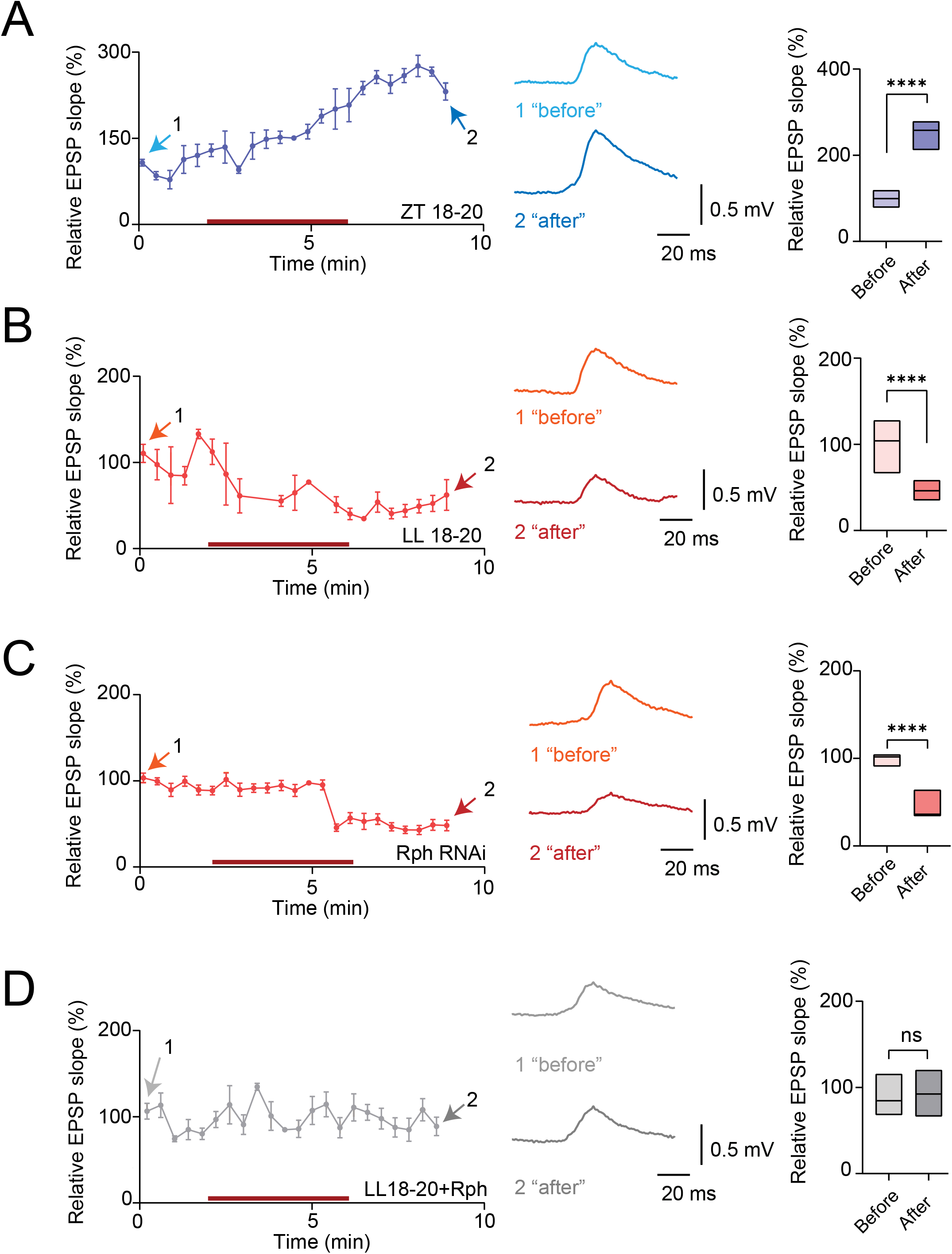
Metaplasticity induced by presynaptic spiking pattern, light, and Rph in the synapses between DN1p and PI neurons. (**A**) Time course of synaptic potentiation observed in PI neurons induced by daytime spiking patterns in DN1p neurons from control flies at ZT18-20 (dark purple). Averaged PSPs recorded before (1, light blue) and after (2, dark blue) stimulation are shown. Quantification of the average relative EPSP slope before (lavender) and after (purple) stimulation is shown at right (N=9). (**B**) Time course of synaptic depression observed in PI neurons induced by the same daytime DN1p spiking patterns paradigm from flies maintained in constant light at LL18- 20 (red-orange). Averaged PSPs recorded before (1, orange) and after (2, dark orange) stimulation are shown. Quantification of the average relative EPSP slope before (pink) and after (red) stimulation is shown at right (N=5). (**C**) Time course of synaptic depression observed in PI neurons induced by the same daytime DN1p spiking patterns paradigm from flies expressing Rph RNAi in DN1p neurons at ZT18-20 (red-orange). Averaged PSPs recorded before (1, orange) and after (2, dark orange) stimulation are shown. Quantification of the average relative EPSP slope before (pink) and after (red) stimulation is shown at right (N=20). (**D**) Time course of synaptic potentials in PI neurons during the same daytime DN1p spiking patterns stimulation from flies maintained in constant light at LL18-20 after synthesized Rph addition (gray). Averaged PSPs recorded before (1, light gray) and after (2, dark gray) stimulation are shown. Quantification of the average relative EPSP slope before (light gray) and after (dark gray) stimulation is shown at right (N=11). The statistics used were an unpaired t-test with *p < 0.05, **p < 0.01, ***p < 0.001, ****p < 0.0001, and ns indicated non-significant.

### Circadian state gates bidirectional timing-dependent plasticity in PI circuits

We next asked whether circadian state also controls the temporal learning rules of the DN1p→PI circuit. Bidirectional pairing experiments performed at night (ZT18–20) revealed timing-dependent plasticity in control flies: one pairing order induced synaptic potentiation (**Fig. 5A**), whereas the opposite order induced synaptic depression (**Fig. 5C**). Thus, the nighttime PI circuit retains the capacity for bidirectional synaptic modification based on temporal contingency. To define the mechanism of the potentiating branch, we reduced NMDA receptor signaling using dsNR1. This manipulation attenuated or abolished pairing-induced potentiation (**Fig. 5B**), consistent with an NMDAR-dependent Hebbian component (46). Constant-light exposure altered the depressive branch of the timing rule (**Fig. 5D**), indicating that circadian perturbation changes not only the basal synaptic state but also the translation of spike timing into lasting synaptic modification. Taken together with the homosynaptic plasticity data, these experiments identify Rph-dependent circadian state as a key determinant of metaplastic thresholds in the DN1p→PI circuit in *Drosophila*.

**Fig. 5.**
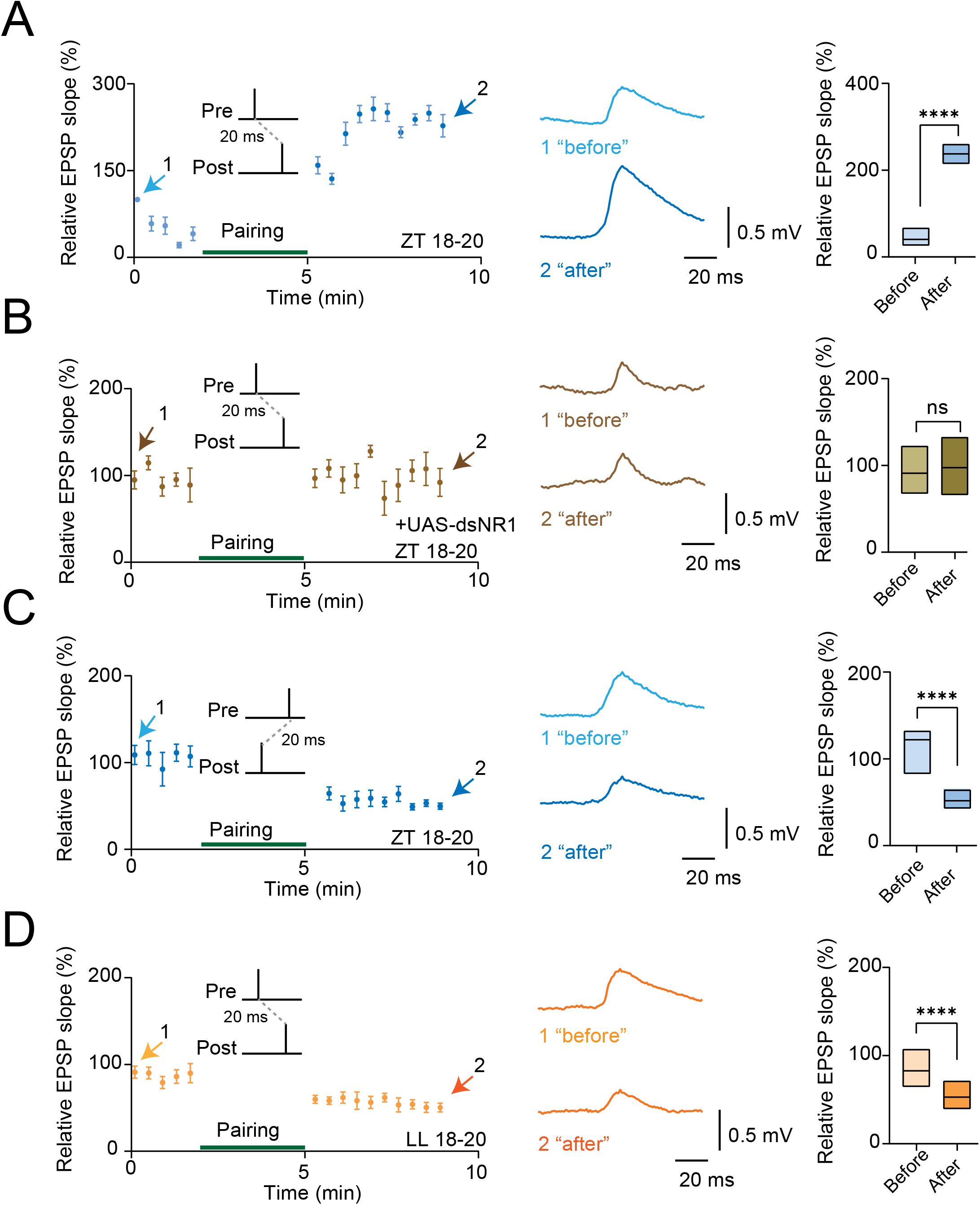
Metaplasticity induced by spike timing and light in the synapses between DN1p and PI neurons. (**A**) Time course of LTP induced by pairing in PI neurons from control flies at ZT18–20. Averaged PSPs before (1, light blue) and after (2, dark blue) pairing are shown. Quantification of the average relative EPSP slope before (light blue) and after (dark blue) pairing is shown at right (N=6). (**B**) Time course of synaptic responses induced by the same pairing paradigm in flies expressing UAS-dsNR1 at ZT18–20. Averaged PSPs before and after pairing are shown. Quantification of the average relative EPSP slope before and after pairing is shown at right (N=10). (**C**) Time course of synaptic depression induced by pairing in PI neurons from control flies at ZT18–20. Averaged PSPs before (1, light blue) and after (2, dark blue) pairing are shown. Quantification of the average relative EPSP slope before (light blue) and after (dark blue) pairing is shown at right (N=10). (**D**) Time course of synaptic depression induced by pairing in PI neurons from flies maintained in constant light at LL18– 20. Averaged PSPs before (1, light orange) and after (2, dark orange) pairing are shown. Quantification of the average relative EPSP slope before (light orange) and after (dark orange) pairing is shown at right (N=15). The statistics used were an unpaired t-test with *p < 0.05, **p < 0.01, ***p < 0.001, ****p < 0.0001, and ns indicated non-significant.

## Discussion

We show that Rabphilin (Rph) as a molecular effector through which circadian time is translated into a metaplastic state in a sleep-regulating circuit in *Drosophila*. Rph levels increase at night, when DN1p- dependent sleep consolidation is most prominent, and Rph is required in DN1p clock neurons for stable nighttime sleep. While we found that Rph has little effect on spike output, it strongly reshapes the temporal structure of subthreshold membrane-voltage fluctuations in DN1p neurons. These changes are transmitted from DN1p presynaptic biophysics to downstream PI neurons as altered synaptic variability, synchrony and plasticity thresholds. Together, the data support a hierarchical model in which a clock-regulated synaptic protein tunes membrane dynamics in pacemaker neurons and thereby sets the metaplastic state of a sleep-regulating circuit. Rph RNAi knockdown left mean spontaneous firing, several measures of spike-train variability and evoked firing largely intact, yet significantly altered the stochastic process of voltage fluctuations in the membrane potential of DN1p neurons. These findings indicate that circadian clock neural encoding mechanisms engineering sleep is embedded not only in action potential output, but also in the stochastic structure of the membrane potential itself. In this framework, subthreshold dynamics are not passive background noise; they shape how incoming and internally generated signals are filtered over time and thus bias the circuit toward distinct functional and plasticity states. The nightly enrichment of Rph places this mechanism within a molecular circadian framework. The apparent dissociation between protein abundance and transcript-level measurements suggests that Rph is regulated at least in part beyond transcription, potentially through rhythmic control of protein trafficking or post-translational modification (47-49). The similarity between Rph depletion and light-induced circadian disruption (50-52) further supports the idea that normal nocturnal accumulation of Rph helps establish a circuit state that favors consolidated sleep, whereas inappropriate light exposure destabilizes this state. At the circuit level, Rph does not simply scale DN1p-to-PI transmission. Instead, it reorganizes the temporal statistics of synaptic signaling, including PSP variance, event timing, state-transition structure and bilateral synchrony. These properties are important because they define the regime in which synaptic modification rules are expressed. Rph therefore acts metaplastically: it changes the conditions under which subsequent plasticity is induced, rather than imposing a fixed increase or decrease in synaptic strength. The ability of synthesized Rph introduction to be able to shift PSP dynamics and bias the direction of synaptic plasticity change under perturbed circadian conditions further argues that Rph is not merely a marker of circuit state, but an active determinant of it. Under dim nocturnal light stimulation, i.e., “light pollution”(53-59), we found that Rph RNAi knockdown permitted synaptic potentiation, whereas exogenous Rph biased the same pathway toward depression. Under optogenetic manipulations that mimic daytime presynaptic DN1ps spiking during nighttime (21), these relationships reversed. Rph therefore does not behave as a constitutive promoter of depression or suppressor of potentiation. Rather, it acts as a bidirectional regulator of the synaptic modification threshold whose effect depends on circadian phase and recent activity. Recruitment of postsynaptic PI neuronal spiking revealed an additional layer of regulation: timing-dependent plasticity emerged, and its polarity was determined by light. Circadian state thus acts upstream of the temporal learning rules themselves (22, 60-67), specifying when a given spike-timing relationship is interpreted as a strengthening or weakening signal (68, 69). These results support a hierarchical organization of stabilizing mechanisms in the sleep circuit. A primary Rph- dependent process appears to constrain baseline synaptic state and limit inappropriate strengthening under ordinary nocturnal conditions (36, 70), whereas timing-dependent plasticity becomes engaged when synapses enter regimes that recruit postsynaptic spiking (71-76). Such an architecture would allow the circuit to remain robust in the face of ongoing perturbation while preserving sufficient flexibility to incorporate salient environmental information. More broadly, the work provides a mechanistic bridge from microscale membrane dynamics to network computation and behavioral state, showing how relatively small changes in structured neural variability can propagate to downstream plasticity and ultimately alter sleep. This framework also reframes circadian regulation of sleep. Sleep is often considered a downstream consequence of relatively static clock output or changes in firing rate within pacemaker neurons (77-79). The present data instead support a model in which the clock continuously tunes the computational state of sleep circuits by modulating membrane restoration dynamics and synaptic plasticity thresholds. Nighttime sleep consolidation may therefore depend less on increasing sleep-promoting drive per se than on configuring a synaptic landscape that filters perturbation, constrains inappropriate potentiation and preserves appropriate sensitivity to salient input (35, 67, 80, 81). In this sense, circadian regulation of sleep emerges as an active metaplastic process that balances robustness and adaptive flexibility across timescales. Several limitations define important directions for future work. Because the present study focuses on the neurophysiological characterization, future studies will need to determine how the altered membrane-fluctuation parameters map onto specific vesicle cycling and metabolic state, and to identify the binding partners and post-translational mechanisms that confer circadian control over Rph function. In addition. light was the principal environmental perturbation examined here; other zeitgebers and/or temperature (82) may reveal additional routes through which circadian circuits tune metaplastic state. Taken together, these findings establish a model in which circadian clock sets a metaplastic setpoint by regulating subthreshold membrane dynamics in *Drosophila*. Through this mechanism, microscale electrical variability is translated into changes in synaptic timing, plasticity polarity and sleep stability, linking molecular timekeeping to circuit computation and behavioral-state control.

## Materials and Methods

### *Drosophila* genetics

All *Drosophila* lines in this study were obtained from the Bloomington *Drosophila* Stock Center (Bloomington, IN, USA), except for *UAS-Rph RNAi* (VDRC: 52438 and 109337) and *UAS-RNAi control* (GFP RNAi) lines (VDRC: 60103) obtained from the Vienna Drosophila Research Center, *UAS-dsNR1 RNAi line* (a gift from C.- L. Wu), and *dilp2-mCherry* (a gift from A. Sehgal). For the sleep experiments shown in **Figs. 1 and S1**, the *R18H11-Gal4* line (BDSC: 48832) was crossed with either *UAS-Rph RNAi* (VDRC: 52438 and 109337) or control *UAS-GFP RNAi* (VDRC: 60103) lines. For electrophysiological experiments in **Figs. 1, 2, and S2**, the *R18H11-Gal4* line (BDSC: 48832) and *UAS-CD4-tdGFP* line (BDSC: 35836) were recombined through standard genetic recombination techniques and this was crossed with *UAS-Rph RNAi* line. For experiments in **Figs. 3, 4, S3, and S4**, flies having *R18H11-Gal4* line (BDSC: 48832), *dilp2-mCherry* (a gift from A. Sehgal) > *UAS-IVS-CsChrimson*.*mVenus* line (BDSC:55135) were crossed with *UAS-Rph RNAi* lines were used. For experiments in **Fig. 5**, the *R18H11-Gal4* line (BDSC: 48832), the *dilp2-Gal4* line (BDSC: 37516) and *UAS-IVS-CsChrimson*.*mVenus* line (BDSC: 55135) were recombined through standard genetic recombination techniques and this was crossed with *UAS-dsNR1 RNAi* line (a gift from C.-L. Wu) was used. Flies were kept in an incubator (DR-36VL, Percival Scientific, Perry, IA, United States) under 12 h:12 h light-dark cycles and 65% humidity. Files were fed standard *Drosophila* food containing molasses, cornmeal, and yeast. Flies used for all experiments were female with the age range of 5-8 days old. All experiments were conducted at 25 °C and were performed in compliance with all relevant ethical regulations for animal testing and research at Case Western Reserve University.

### Electrophysiology

Cold-anesthetized flies were immobilized on metal shims (0.025 mm) using dental wax or UV-curable adhesive. Brain exposure involved cuticle removal and placement in carbogen-saturated saline (101 mM NaCl, 3 mM KCl, 1 mM CaCl_2_, 4 mM MgCl_2_, 1.25 mM NaH_2_PO_4_, 20.7 mM NaHCO_3_, 5 mM glucose; 235-245 mOsm, pH 7.2). Enzymatic treatment (collagenase 0.1 mg/mL, protease XIV 0.2 mg/mL, dispase 0.3 mg/mL) for 60 seconds facilitated glial removal. Neurons were identified through fluorescence (CoolLED PE4000) on an Olympus BX51WI microscope. To minimize access resistance variability inherent in conventional glass electrodes, we implemented membrane-coated electrode techniques (83, 84). The coating material was synthesized by ultrasonicating egg yolk lecithin (7.6 g/L; Sigma-Aldrich 440154) with cholesterol (2 mM; Sigma-Aldrich C8667) in a hexane-acetone mixture for 60 minutes at ambient temperature. Following nitrogen- assisted evaporation and vacuum treatment, the lipid components were resuspended in paraffin-squalene solution (7:3 ratio) and heated overnight at 80°C. Sharp electrodes were pulled from quartz glass capillaries with a filament (outer/inner diameter: 1.2/0.6 mm) using a laser-based puller (P-2000, Sutter Instrument), backfilled with 1 M KCl, and had resistances ranging from 120 to 190 MΩ (45, 84). All solutions were filtered through 0.02 μm syringe filters (Anotop 10, Whatman). The electrode subsequently received lipid coating through vertical dipping methodology, where tips contacted internal solution before lipid application (<20 μL), followed by controlled vertical withdrawal. Sharp recordings employed 1 M KCl in the electrodes. For the experiments to test synthesized Rph introduction, recombinant *Drosophila* Rabphilin protein (synthesized by Leading Biology) was introduced via intracellular perfusion. Purified Rabphilin (5 μg/ml final concentration) was added to the internal solution before recording and continuously perfused through electrode holder having perfusion ports (Catalog #:663010, A-M systems). We waited 10–15 minutes after inserting the electrodes to allow for adequate diffusion of intracellular proteins before initiating the plasticity protocols. Sharp recordings targeted axonal regions identified by concentrated fluorescence. Electrode penetration was guided acoustically (A-M Systems 3300) and electrically, with pulse durations optimized at 2-5 ms. Axoclamp 2B amplifiers provided signal acquisition, digitized at 10 kHz through Digidata 1550B with 1 kHz filtering. Optogenetic manipulation of DN1p presynaptic spiking patterns was performed essentially as previously described (21). Flies were fed 1 mM all-trans-retinal (MilliporeSigma) mixed into rehydrated food flakes (Nutri-Fly Instant; Genesee Scientific). Photostimulation of CsChrimson-expressing DN1p neurons was delivered using a collimated 625 nm LED light source (Thorlabs). A fiber optic cannula (Thorlabs) was coupled to the LED to enable focused light delivery. Stimulus pulse patterns, designed to mimic DN1p presynaptic spiking dynamics, were generated based on a Gaussian mixture model (GMM) and delivered via TTL pulses from a dual-channel waveform generator (DigiStim-2; ALA Scientific). Given our evidence that DN1p neurons are coupled via gap junctions (21), the synchrony of DN1p firing under physiological conditions is expected to approximate that induced by optogenetic stimulation. However, we cannot exclude the possibility that optogenetic activation produces a higher degree of synchrony than occurs naturally. Spike timing-dependent plasticity was induced using paired pre- and postsynaptic stimulation protocols. DN1p neurons were stimulated to evoke single action potentials using brief (5 ms) optogenetic stimulations (Optopatcher, A-M systems) through the glass electrodes. Postsynaptic spikes in PI neurons were similarly evoked by using the same optogenetic stimulations before or after presynaptic stimulation. Each pairing consisted of 30 repetitions at 0.2 Hz. For pre-before-post protocols, DN1 stimulation preceded PI stimulation by Δt = +20 ms, while post-before-pre protocols used Δt = -20 ms. Changes in synaptic plasticity in PI neurons were plotted by EPSP slope, but the statistical quantification before and after pairing protocols was only made by using the first and last bin of EPSP slope.

### Ornstein–Uhlenbeck (OU) model

An Ornstein–Uhlenbeck (OU) model was built using parameters derived from membrane voltage recordings. The signals were first preprocessed and mean-centered. The OU time constant was estimated from the temporal autocorrelation by fitting an exponential decay, representing the inverse of the mean-reversion rate. Noise amplitude was measured as the standard deviation of detrended voltage fluctuations. Simulations were performed in the Brian simulator using a fixed time step matching the original sampling interval. The simulated OU signals were analyzed with continuous wavelet transform to separate frequency-specific components. Each frequency band was then tested for stationarity using the Augmented Dickey-Fuller (ADF) test, which evaluates whether a time series contains a unit root. Higher p-values indicate weaker evidence against non-stationarity. Differences in p-value distributions between genotypes were interpreted as differences in the extent of non-stationary dynamics in the OU-modeled signals.

### Sleep Analysis

Sleep behavior was measured with the use of consolidated locomotor inactivity, as described previously (84). Female flies (5-10 days old) were loaded into glass tubes containing 5% sucrose and 2% *Drosophila* agar, for sleep analysis using the *Drosophila* Activity Monitor (DAM) from Trikinetics. Flies were exposed to two days of 12 hr light/12 hr dark followed by 24 hr darkness for five days. Monitors were read every 30 seconds, and data was analyzed using MATLAB. The first day following loading was not included in the analysis. Activity counts were collected in 1 min bins, and sleep was identified as periods of inactivity of at least 5 min. Sleeplab (85), a MATLAB-based (MathWorks) software, was used to quantify sleep behavior. Single-fly video analyses were conducted as previously described (21). Flies were anesthetized on ice and affixed to a 0.025 mm–thick stainless steel shim using dental wax. Flies were provided ad libitum access to food. Leg movements were continuously recorded using an infrared (IR)-sensitive CCD camera at 2 frames per second. Inactivity, inferred from leg movement, was converted into sleep behavior using a frame-subtraction method with a noise-threshold algorithm(21). Micromovement was quantified by identifying “010” sequences (1 = active, 0 = inactive) in 1- minute binned data. An “active” bin was defined as any instance in which pixel intensity changes exceeded a 2 standard deviation threshold. A total of 90 minutes of recording was used for analysis, as the first 30 minutes of the 2-hour recording period were excluded to allow flies to acclimate to the experimental setup.

### qPCR

To quantify Rph transcripts in DN1ps, quantitative PCR (qPCR) was performed by using harvested DN1ps. Fly brains were dissected in ice-cold dissecting saline and collected in cold saline throughout the procedure. Samples were centrifuged at room temperature for 2 min at 1,000 rpm, after which the supernatant was removed and brains were washed in 300 μl of cold dissecting saline. An enzyme cocktail was added at a volume of approximately 2 μl per brain, and samples were incubated at room temperature for 20–30 min with occasional gentle mixing using a 20-μl pipette tip. Enzymatic digestion was quenched by adding a fivefold volume of cold dissecting saline. Brains were then centrifuged for 2 min at room temperature and resuspended in cold SM- active Bis–Tris medium at 7 μl per brain, with a minimum total volume of 400 μl to facilitate trituration. Mechanical dissociation was performed by sequential trituration using flame-rounded glass pipette tips of decreasing diameter: 30 passes with a large-bore tip, 20 passes with a medium-bore tip, and 10 passes with a small-bore tip. Samples were placed on ice after every 10 pipetting cycles to maintain low temperature. Trituration was continued until the suspension flowed freely through the small-bore tip, taking care to avoid over-trituration, which can result in cell lysis. Following dissociation, brains were transferred to a dish and visually identified under a fluorescence stereomicroscope. Individual fluorescently labeled DN1ps neurons were isolated using glass micropipettes pulled from capillary tubes with a micropipette puller. Prior to use, micropipette tips were gently broken to achieve an opening approximately the width of a single cell. Target DN1ps cells were approached under visual guidance and aspirated into the micropipette using gentle mouth suction. The micropipette was then transferred to a tube containing 100 μl of TRIzol reagent (Invitrogen), and the tip was gently broken to release the collected DN1ps cells. 100 DN1ps neurons were harvested per sample into a 0.2-ml tube containing 100 μl TRIzol and immediately frozen at −80 °C until further processing. Total RNA was extracted from these harvested DN1ps neurons using TRIzol Reagent according to the manufacturer’s instructions. qPCR was carried out using SYBR PCR Master Mix (Applied Biosystems) on a 7900 Real-Time PCR System (Applied Biosystems). Primers targeting Rph were used as follows: Rph-RA F: 5’- CCCTTCGTCAAGATCCAACTC-3’ and Rph-RA-R: 5’- CTCCTCGTTGTAGATGGGATTG-3’. RpL32 was used as an internal control. Ct values were quantified by comparison to a regression-based standard curve generated using the same primer pairs. Three biological replicates were analyzed for each condition.

### Western Blotting

Samples of *Drosophila* heads were collected and frozen in liquid nitrogen, then emulsified into extraction buffer containing Pierce protease inhibitor tab (Thermo Fisher, A32963), NuPAGE LDS Sample buffer (Fisher Sci, NP0007), NuPAGE sample reducing agent (Fisher Sci, NP0004), and Pierce DTT (Thermo Fisher, 20290). The samples were then boiled at 95°C for 5 minutes, followed by centrifugation for 3-5 minutes at 15,000 rpm. The supernatants were transferred to fresh Eppendorf tubes and protein content was measured using a Nanodrop Spectrophotometer. Samples were loaded into Invitrogen mini pre-cast gels for gel electrophoresis at 100 V for approximately 2 hours. The gel was taken apart and transferred to a nitrocellulose membrane using the Invitrogen mini blot module (Thermo Fisher) at 30 V for 1 hour. After blocking in 5% NGS-PBST for 2 hours, the membrane was left in rabbit anti-Rabphilin-3a (Sigma-Aldrich, 1:1,000, Cat# R3026, RRID:AB_1079836) overnight at 4°C. The membrane was then washed in PBST for at least 15 minutes and placed in goat anti-rabbit HRP (Sigma, 1:10,000) for 1 hour at room temperature. The HRP was activated using Pierce ECL Substrate (Thermo Fisher) and left for 10 minutes to incubate. The membrane was imaged using a ChemiDoc imaging system (BioRad) in chemiluminescent mode. Band intensity was determined using ImageJ.

### Immunohistochemistry

*Drosophila* brains were fixed in 4% PFA for 30 min at room temperature. After several washes with phosphate-buffered saline (137 mM NaCl, 2.7 mM KCl, 10 mM Na_2_HPO_4_, 1.7 mM KH_2_PO_4_) + 0.3% Triton X-100 (PBST), samples were incubated with chicken anti-GFP (Thermo Fisher, 1:200, Cat# A10262; RRID: AB_2534023) and rabbit anti-Rabphilin-3a (Rockland, 1:1,000, Cat# 612-401-E21, RRID:AB_11182467) at 4°C overnight. After additional PBST washes, samples were incubated with Alexa Fluor 488 Goat anti-Chicken IgY (Thermo Fisher, 1:1000, Cat# A11039; RRID: AB_2534096) for anti-GFP staining and Alexa Fluor 568 Goat anti-Rabbit IgG (Thermo Fisher, 1:1000, Cat# A11011; RRID: AB_143157) for anti-Rph stainings overnight at 4°C. After another series of washes in PBST at room temperature over 1 hr, samples were cleared in 70% glycerol in PBS for 5 min at room temperature and then mounted in Vectashield (Vector Labs). Confocal microscope images were taken under 10x or 63x magnification using a Zeiss LSM800.

### Statistical Methods

All statistical and data analyses were performed using Prism version 10.6.1 (GraphPad), Clampfit version 10.7 (Molecular Devices), and MATLAB R2025b (MathWorks). For statistical analysis, two-group comparisons were conducted using unpaired t-tests. When comparing multiple groups, one-way ANOVA followed by Tukey’s post-hoc test was applied to normally distributed datasets, whereas the Kruskal-Wallis test with Dunn’s multiple comparisons test was used for non-normally distributed datasets. Differences were considered statistically significant at p < 0.05. Asterisks indicate significance levels: *p < 0.05, **p < 0.01, ***p < 0.001, and ****p < 0.0001. Error bars represent the standard error of the mean (SEM), averaged across all experiments.

## Acknowledgments

We thank Yoji Yamamoto for initial processing of the sleep dataset from our large-scale RNAi screen, and Ayaka Fujimaki and Yusuke Kurisu for technical assistance with Ornstein–Uhlenbeck (OU) computational modeling. We also thank Ben Strowbridge, Heather Broihier, Dominique Durand, and members of the Tabuchi lab for helpful discussions. This work was supported by grants from the National Institutes of Health (R00NS101065 and R35GM142490), Whitehall Foundation, BrightFocus Foundation (A2021043S), PRESTO grant from Japan Science and Technology Agency (JPMJPR2386), Research Corporation for Science Advancement (SA-MBC-2024-080c), and the Tomizawa Jun-ichi and Keiko Fund of the Molecular Biology Society of Japan for Young Scientists.

## Figure Legends

**Supplementary Fig. 1.** Additional sleep phenotypes and immunostaining of Rph in DN1p neurons.

(**A**) Twenty-four-hour sleep profiles of *R18H11-GAL4* flies (gray, N = 32), *R18H11-GAL4>UAS-Rph RNAi #1* flies (green, N = 32), and *R18H11-GAL4>UAS-Rph RNAi #2* flies (blue, N = 32). (**B–D**) Quantification of sleep parameters during ZT6–8 and ZT18–20 for the genotypes shown in **A**. (**B**) Sleep-bout duration. (**C**) Sleep-bout number. (**D**) Total sleep time. (**E)** Immunohistochemistry showing Rph in DN1p cell bodies using anti-Rph (Rabphilin 3A) (magenta) and anti-GFP (green) labeling. **p < 0.01, ***p < 0.001, ****p < 0.0001. ns: non-significance based on one-way ANOVA followed by post-hoc Tukey tests.

**Supplementary Fig. 2.** Additional electrophysiological analyses of DN1p firing and intrinsic membrane properties assessed by current injection.

(**A**) Representative membrane-potential traces showing current-evoked firing in control flies (gray) and flies expressing Rph RNAi in DN1p neurons (orange). (**B**) Firing rate–current (*f–I*) relationship for control flies (N = 8) and flies expressing Rph RNAi in DN1p neurons (N = 6) at ZT18-20; slope factors from linear regression are compared. (**C**) First-spike latency as a function of injected current in control flies and flies expressing Rph RNAi in DN1p neurons at ZT18-20. (**D**) Boltzmann fits to the time course of reciprocal interspike interval (1/ISI) during current-evoked firing in control flies and flies expressing Rph RNAi in DN1p neurons at ZT18- 20. (**E**) Half-time derived from the fits shown in **D** (control, N = 20; Rph RNAi, N = 12). (**F**) Plateau-arrival time derived from the fits shown in **D** (control, N = 20; Rph RNAi, N = 12). (G) Input resistance measured from membrane-voltage responses to current injection at ZT18-20 (control, N = 32; Rph RNAi, N = 17). (**H**) Distribution of spike-onset velocity values from spontaneous action potentials, with superimposed example waveforms for control flies and flies expressing Rph RNAi in DN1p neurons at ZT18-20. (**I**) Phase plots of action potential dynamics (dV/dt versus membrane potential) from spontaneous firing in control flies and flies expressing Rph RNAi in DN1p neurons at ZT18-20. The statistics used were an unpaired t-test with *p < 0.05, **p < 0.01, ***p < 0.001, ****p < 0.0001, and ns indicated non-significant.

**Supplementary Fig. 3.** Additional electrophysiological analyses of spontaneous PSPs in PI neurons.

(**A**) Superimposed traces of spontaneous PSPs recorded from PI neurons in control flies (gray) at ZT18-20 (N=8), flies maintained in constant light (yellow) at LL18-20 (N=8), and flies expressing Rph RNAi in DN1p neurons (orange) at ZT18-20 (N=8). Representative miniature PSPs (mPSPs) and large PSPs are shown for each condition. (**B**) Variability of mPSP amplitudes in PI neurons. (**C**) Cumulative distributions of mPSP amplitude. (**D**) mPSP slope. (**E**) Histogram of inter-mPSP interval distributions. (**F**) Quantification of inter-mPSP intervals. **p < 0.01, ***p < 0.001, ****p < 0.0001. ns: non-significance based on one-way ANOVA followed by post-hoc Tukey tests.

**Supplementary Fig. 4.** Light/clock-generated neural computations controlling circadian regulation of sleep in the synapses between DN1p and PI neurons.

(**A**) Representative examples of near-synchronous PSPs in dual-electrode recordings from PI neurons in flies maintained in constant light (yellow) at LL18-20 and flies expressing Rph RNAi in DN1p neurons (orange) at ZT18-20. (**B**) Schematics summarizing the main conclusions of the study. Environmental light shifts the synaptic setpoint toward a potentiation-prone, unstable DN1p–PI state associated with reduced sleep quality, whereas the circadian clock, through Rph, biases the setpoint toward a depression-prone, stable state associated with improved sleep quality. This metaplastic setpoint regulates sleep–wake behavior and influences the direction of plasticity at DN1p–PI synapses. When postsynaptic spiking occurs, these synapses also exhibit spike-timing-dependent plasticity (STDP), with Hebbian potentiation and anti-Hebbian depression. The coexistence of STDP and previously identified “spike pattern-dependent plasticity (SPDP)” (21) suggests multiplexed temporal coding mechanisms based on timing and patterns information that may shape arousal and sleep in *Drosophila*.

## References

1. C. I. Bargmann, Beyond the connectome: how neuromodulators shape neural circuits. Bioessays 34, 458–465 (2012).

2. S. J. Martin, P. D. Grimwood, R. G. Morris, Synaptic plasticity and memory: an evaluation of the hypothesis. Annu Rev Neurosci 23, 649–711 (2000).

3. E. Marder, J. M. Goaillard, Variability, compensation and homeostasis in neuron and network function. Nat Rev Neurosci 7, 563–574 (2006).

4. E. Marder, Neuromodulation of neuronal circuits: back to the future. Neuron 76, 1–11 (2012).

5. S. Grillner, The motor infrastructure: from ion channels to neuronal networks. Nat Rev Neurosci 4, 573–586 (2003).

6. C. Dubowy, A. Sehgal, Circadian Rhythms and Sleep in Drosophila melanogaster. Genetics 205, 1373–1397 (2017).

7. P. E. Hardin, S. Panda, Circadian timekeeping and output mechanisms in animals. Curr Opin Neurobiol 23, 724–731 (2013).

8. R. Allada, B. Y. Chung, Circadian organization of behavior and physiology in Drosophila. Annu Rev Physiol 72, 605–624 (2010).

9. H. D. Piggins, A. Loudon, Circadian biology: clocks within clocks. Curr Biol 15, R455–457 (2005).

10. J. Brankack, V. I. Kukushka, A. L. Vyssotski, A. Draguhn, EEG gamma frequency and sleep-wake scoring in mice: comparing two types of supervised classifiers. Brain Res 1322, 59–71 (2010).

11. J. Cedernaes, N. Waldeck, J. Bass, Neurogenetic basis for circadian regulation of metabolism by the hypothalamus. Genes Dev 33, 1136–1158 (2019).

12. C. S. Colwell, Linking neural activity and molecular oscillations in the SCN. Nat Rev Neurosci 12, 553–569 (2011).

13. L. Zhang et al., DN1(p) circadian neurons coordinate acute light and PDF inputs to produce robust daily behavior in Drosophila. Curr Biol 20, 591–599 (2010).

14. A. Seluzicki et al., Dual PDF signaling pathways reset clocks via TIMELESS and acutely excite target neurons to control circadian behavior. PLoS Biol 12, e1001810 (2014).

15. M. Flourakis et al., A Conserved Bicycle Model for Circadian Clock Control of Membrane Excitability. Cell 162, 836–848 (2015).

16. A. Lamaze, R. Stanewsky, DN1p or the “Fluffy” Cerberus of Clock Outputs. Front Physiol 10, 1540 (2019).

17. F. Guo et al., Circadian neuron feedback controls the Drosophila sleep--activity profile. Nature 536, 292–297 (2016).

18. F. Guo, M. Holla, M. M. Diaz, M. Rosbash, A Circadian Output Circuit Controls Sleep-Wake Arousal in Drosophila. Neuron 100, 624–635 e624 (2018).

19. A. Lamaze, P. Kratschmer, K. F. Chen, S. Lowe, J. E. C. Jepson, A Wake-Promoting Circadian Output Circuit in Drosophila. Curr Biol 28, 3098–3105 e3093 (2018).

20. M. N. Nitabach, P. H. Taghert, Organization of the Drosophila circadian control circuit. Curr Biol 18, R84–93 (2008).

21. M. Tabuchi et al., Clock-Generated Temporal Codes Determine Synaptic Plasticity to Control Sleep. Cell 175, 1213–1227 e1218 (2018).

22. M. Tabuchi, K. E. Coates, O. B. Bautista, L. H. Zukowski, Light/Clock Influences Membrane Potential Dynamics to Regulate Sleep States. Front Neurol 12, 625369 (2021).

23. Y. Zhang, Y. Liu, D. Bilodeau-Wentworth, P. E. Hardin, P. Emery, Light and temperature control the contribution of specific DN1 neurons to Drosophila circadian behavior. Curr Biol 20, 600–605 (2010).

24. A. Sehgal et al., Molecular analysis of sleep: wake cycles in Drosophila. Cold Spring Harb Symp Quant Biol 72, 557–564 (2007).

25. A. Sehgal et al., Rhythmic expression of timeless: a basis for promoting circadian cycles in period gene autoregulation. Science 270, 808–810 (1995).

26. P. E. Hardin, J. C. Hall, M. Rosbash, Feedback of the Drosophila period gene product on circadian cycling of its messenger RNA levels. Nature 343, 536–540 (1990).

27. R. Allada, N. E. White, W. V. So, J. C. Hall, M. Rosbash, A mutant Drosophila homolog of mammalian Clock disrupts circadian rhythms and transcription of period and timeless. Cell 93, 791–804 (1998).

28. X. Chen, M. Rosbash, MicroRNA-92a is a circadian modulator of neuronal excitability in Drosophila. Nat Commun 8, 14707 (2017).

29. S. Liu et al., WIDE AWAKE mediates the circadian timing of sleep onset. Neuron 82, 151–166 (2014).

30. C. N. Allen, M. N. Nitabach, C. S. Colwell, Membrane Currents, Gene Expression, and Circadian Clocks. Cold Spring Harb Perspect Biol 9 (2017).

31. M. Tabuchi, Dynamic neuronal instability generates synaptic plasticity and behavior: Insights from Drosophila sleep. Neurosci Res 198, 1–7 (2024).

32. A. Neishabouri, A. A. Faisal, Axonal noise as a source of synaptic variability. PLoS Comput Biol 10, e1003615 (2014).

33. G. B. Ermentrout, R. F. Galan, N. N. Urban, Reliability, synchrony and noise. Trends Neurosci 31, 428–434 (2008).

34. J. Cafaro, F. Rieke, Noise correlations improve response fidelity and stimulus encoding. Nature 468, 964–967 (2010).

35. G. Tononi, C. Cirelli, Sleep and the price of plasticity: from synaptic and cellular homeostasis to memory consolidation and integration. Neuron 81, 12–34 (2014).

36. F. Deak et al., Rabphilin regulates SNARE-dependent re-priming of synaptic vesicles for fusion. EMBO J 25, 2856–2866 (2006).

37. M. Geppert, Y. Goda, C. F. Stevens, T. C. Sudhof, The small GTP-binding protein Rab3A regulates a late step in synaptic vesicle fusion. Nature 387, 810–814 (1997).

38. C. Li et al., Synaptic targeting of rabphilin-3A, a synaptic vesicle Ca2+/phospholipid-binding protein, depends on rab3A/3C. Neuron 13, 885–898 (1994).

39. S. Schoch et al., RIM1alpha forms a protein scaffold for regulating neurotransmitter release at the active zone. Nature 415, 321–326 (2002).

40. M. Geppert, T. C. Sudhof, RAB3 and synaptotagmin: the yin and yang of synaptic membrane fusion. Annu Rev Neurosci 21, 75–95 (1998).

41. D. A. Rusakov, L. P. Savtchenko, P. E. Latham, Noisy Synaptic Conductance: Bug or a Feature? Trends Neurosci 43, 363–372 (2020).

42. G. Tononi, C. Cirelli, Sleep function and synaptic homeostasis. Sleep Med Rev 10, 49–62 (2006).

43. C. Ferrer-Orta et al., Structural characterization of the Rabphilin-3A-SNAP25 interaction. Proc Natl Acad Sci U S A 114, E5343–E5351 (2017).

44. J. Ubach, J. Garcia, M. P. Nittler, T. C. Sudhof, J. Rizo, Structure of the Janus-faced C2B domain of rabphilin. Nat Cell Biol 1, 106–112 (1999).

45. D. L. Nguyen et al., Age-Related Unstructured Spike Patterns and Molecular Localization in Drosophila Circadian Neurons. Front Physiol 13, 845236 (2022).

46. A. Gonzalez-Rueda, V. Pedrosa, R. C. Feord, C. Clopath, O. Paulsen, Activity-Dependent Downscaling of Subthreshold Synaptic Inputs during Slow-Wave-Sleep-like Activity In Vivo. Neuron 97, 1244–1252 e1245 (2018).

47. A. Mehra, C. L. Baker, J. J. Loros, J. C. Dunlap, Post-translational modifications in circadian rhythms. Trends Biochem Sci 34, 483–490 (2009).

48. C. C. Yeung et al., Circadian regulation of protein cargo in extracellular vesicles. Sci Adv 8, eabc9061 (2022).

49. S. Hegazi et al., UBR4/POE facilitates secretory trafficking to maintain circadian clock synchrony. Nat Commun 13, 1594 (2022).

50. R. Lee, A. Tapia, S. Kaladchibachi, M. A. Grandner, F. X. Fernandez, Meta-analysis of light and circadian timekeeping in rodents. Neurosci Biobehav Rev 123, 215–229 (2021).

51. S. Kaladchibachi, D. C. Negelspach, J. M. Zeitzer, F. X. Fernandez, A millisecond parameter space for phase-shifting the circadian pacemaker with near-ultraviolet light. J Comp Physiol A Neuroethol Sens Neural Behav Physiol 211, 551–560 (2025).

52. Y. Wang, K. N. Paul, G. D. Block, T. Deboer, C. S. Colwell, Dim light at night disrupts the sleep-wake cycle and exacerbates abnormal EEG activity in Cntnap2 knockout mice: implications for autism spectrum disorders. Mol Autism 16, 62 (2025).

53. L. K. Fonken, R. J. Nelson, The effects of light at night on circadian clocks and metabolism. Endocr Rev 35, 648–670 (2014).

54. L. K. Fonken, T. G. Aubrecht, O. H. Melendez-Fernandez, Z. M. Weil, R. J. Nelson, Dim light at night disrupts molecular circadian rhythms and increases body weight. J Biol Rhythms 28, 262–271 (2013).

55. L. Tahkamo, T. Partonen, A. K. Pesonen, Systematic review of light exposure impact on human circadian rhythm. Chronobiol Int 36, 151–170 (2019).

56. M. G. Figueiro, Disruption of Circadian Rhythms by Light During Day and Night. Curr Sleep Med Rep 3, 76–84 (2017).

57. R. J. Reiter et al., Light at night, chronodisruption, melatonin suppression, and cancer risk: a review. Crit Rev Oncog 13, 303–328 (2007).

58. S. K. E. Tam et al., Dim light in the evening causes coordinated realignment of circadian rhythms, sleep, and short-term memory. Proc Natl Acad Sci U S A 118 (2021).

59. K. J. Navara, R. J. Nelson, The dark side of light at night: physiological, epidemiological, and ecological consequences. J Pineal Res 43, 215–224 (2007).

60. J. G. Flyer-Adams et al., Regulation of Olfactory Associative Memory by the Circadian Clock Output Signal Pigment-Dispersing Factor (PDF). J Neurosci 40, 9066–9077 (2020).

61. N. F. Ruby et al., Hippocampal-dependent learning requires a functional circadian system. Proc Natl Acad Sci U S A 105, 15593–15598 (2008).

62. S. Inami, T. Sakai, Circadian photoreceptors are required for light-dependent maintenance of long-term memory in Drosophila. Neurosci Res 185, 62–66 (2022).

63. S. Inami et al., Environmental Light Is Required for Maintenance of Long-Term Memory in Drosophila. J Neurosci 40, 1427–1439 (2020).

64. D. H. Loh et al., Rapid changes in the light/dark cycle disrupt memory of conditioned fear in mice. PLoS One 5 (2010).

65. L. M. Wang, N. A. Suthana, D. Chaudhury, D. R. Weaver, C. S. Colwell, Melatonin inhibits hippocampal long-term potentiation. Eur J Neurosci 22, 2231–2237 (2005).

66. D. Chaudhury, L. M. Wang, C. S. Colwell, Circadian regulation of hippocampal long-term potentiation. J Biol Rhythms 20, 225–236 (2005).

67. F. Fernandez et al., Circadian rhythm. Dysrhythmia in the suprachiasmatic nucleus inhibits memory processing. Science 346, 854–857 (2014).

68. H. Markram, J. Lubke, M. Frotscher, B. Sakmann, Regulation of synaptic efficacy by coincidence of postsynaptic APs and EPSPs. Science 275, 213–215 (1997).

69. Y. X. Fu et al., Temporal specificity in the cortical plasticity of visual space representation. Science 296, 1999–2003 (2002).

70. A. Abramian et al., Rabphilin-3A negatively regulates neuropeptide release, through its SNAP25 interaction. Elife 13 (2024).

71. M. Anisimova et al., Spike-timing-dependent plasticity rewards synchrony rather than causality. Cereb Cortex 33, 23–34 (2022).

72. R. C. Froemke, Y. Dan, Spike-timing-dependent synaptic modification induced by natural spike trains. Nature 416, 433–438 (2002).

73. R. C. Froemke, D. Debanne, G. Q. Bi, Temporal modulation of spike-timing-dependent plasticity. Front Synaptic Neurosci 2, 19 (2010).

74. Y. Dan, M. M. Poo, Spike timing-dependent plasticity: from synapse to perception. Physiol Rev 86, 1033–1048 (2006).

75. H. Markram, W. Gerstner, P. J. Sjostrom, Spike-timing-dependent plasticity: a comprehensive overview. Front Synaptic Neurosci 4, 2 (2012).

76. S. Cassenaer, G. Laurent, Hebbian STDP in mushroom bodies facilitates the synchronous flow of olfactory information in locusts. Nature 448, 709–713 (2007).

77. R. E. A. Sanchez, F. Kalume, H. O. de la Iglesia, Sleep timing and the circadian clock in mammals: Past, present and the road ahead. Semin Cell Dev Biol 126, 3–14 (2022).

78. J. R. Jones, M. C. Tackenberg, D. G. McMahon, Manipulating circadian clock neuron firing rate resets molecular circadian rhythms and behavior. Nat Neurosci 18, 373–375 (2015).

79. C. Mazuski et al., Entrainment of Circadian Rhythms Depends on Firing Rates and Neuropeptide Release of VIP SCN Neurons. Neuron 99, 555–563 e555 (2018).

80. G. Wang, B. Grone, D. Colas, L. Appelbaum, P. Mourrain, Synaptic plasticity in sleep: learning, homeostasis and disease. Trends Neurosci 34, 452–463 (2011).

81. C. Cirelli, Sleep and synaptic changes. Curr Opin Neurobiol 23, 841–846 (2013).

82. Z. Liu et al., Behavioral adaptation to warm conditions via Lim1-mediated acceleration of neuronal clocks. Nat Neurosci 29, 374–386 (2026).

83. A. T. Jameson, L. K. Spera, D. L. Nguyen, E. M. Paul, M. Tabuchi, Membrane-coated glass electrodes for stable, low-noise electrophysiology recordings in Drosophila central neurons. J Neurosci Methods 404, 110079 (2024).

84. B. Chong et al., Neuropeptide-Dependent Spike Time Precision and Plasticity in Circadian Output Neurons. Eur J Neurosci 61, e70037 (2025).

85. W. J. Joiner, A. Crocker, B. H. White, A. Sehgal, Sleep in Drosophila is regulated by adult mushroom bodies. Nature 441, 757–760 (2006).

