## Supplementary figures and images for "Metaplastic sleep regulation in *Drosophila* determined by microscale circadian neural dynamics"

### Supplemental Data 1

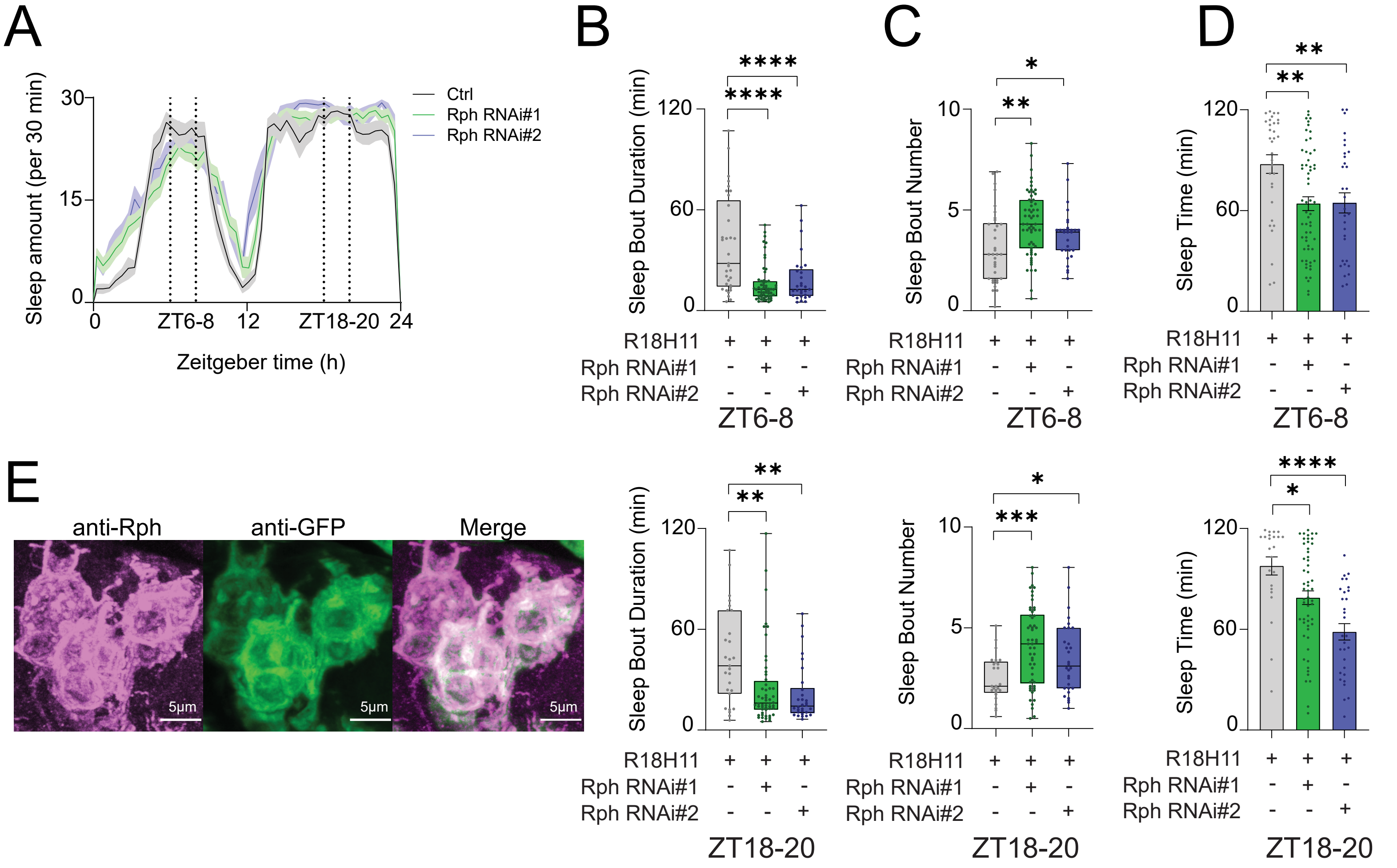

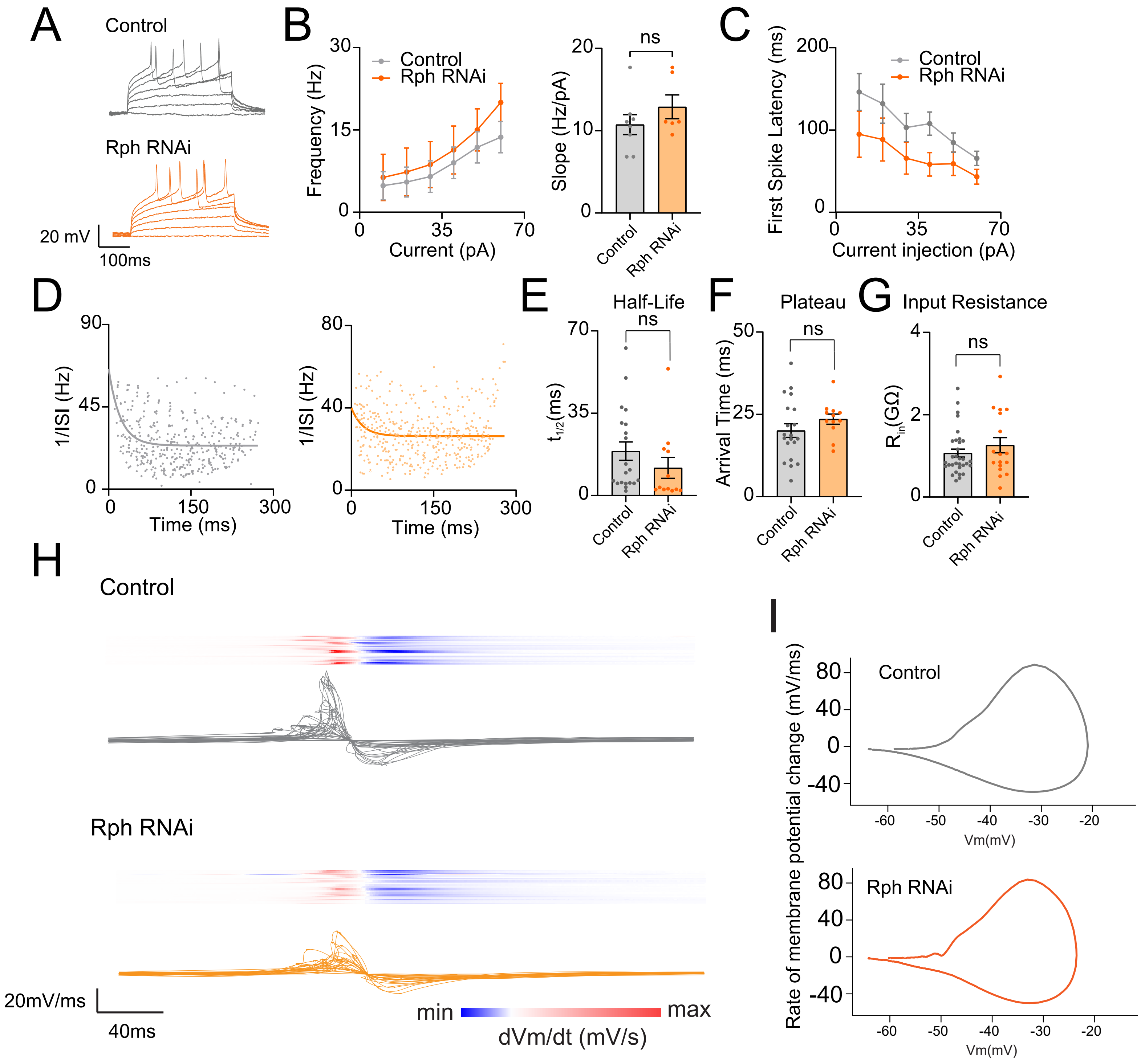

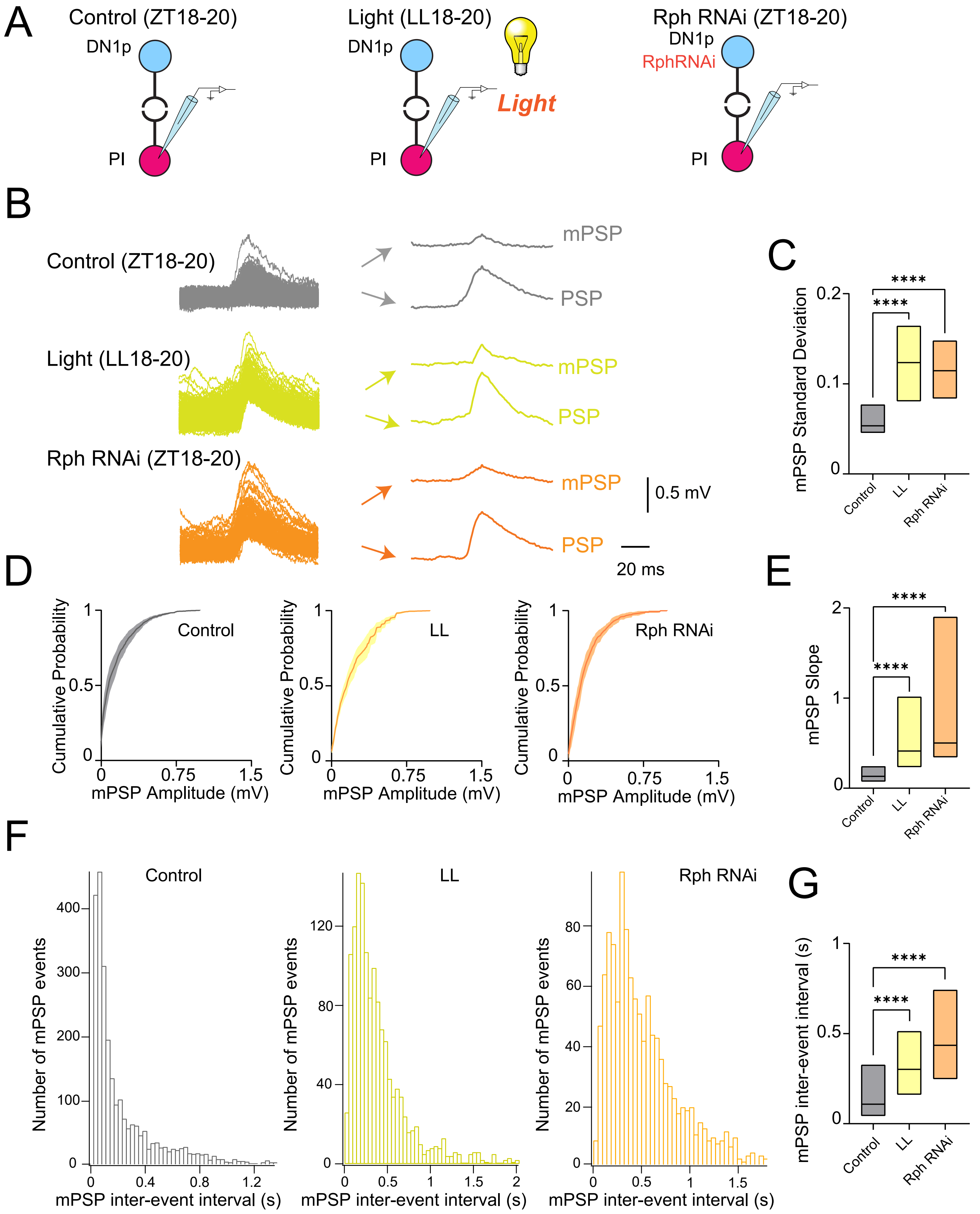

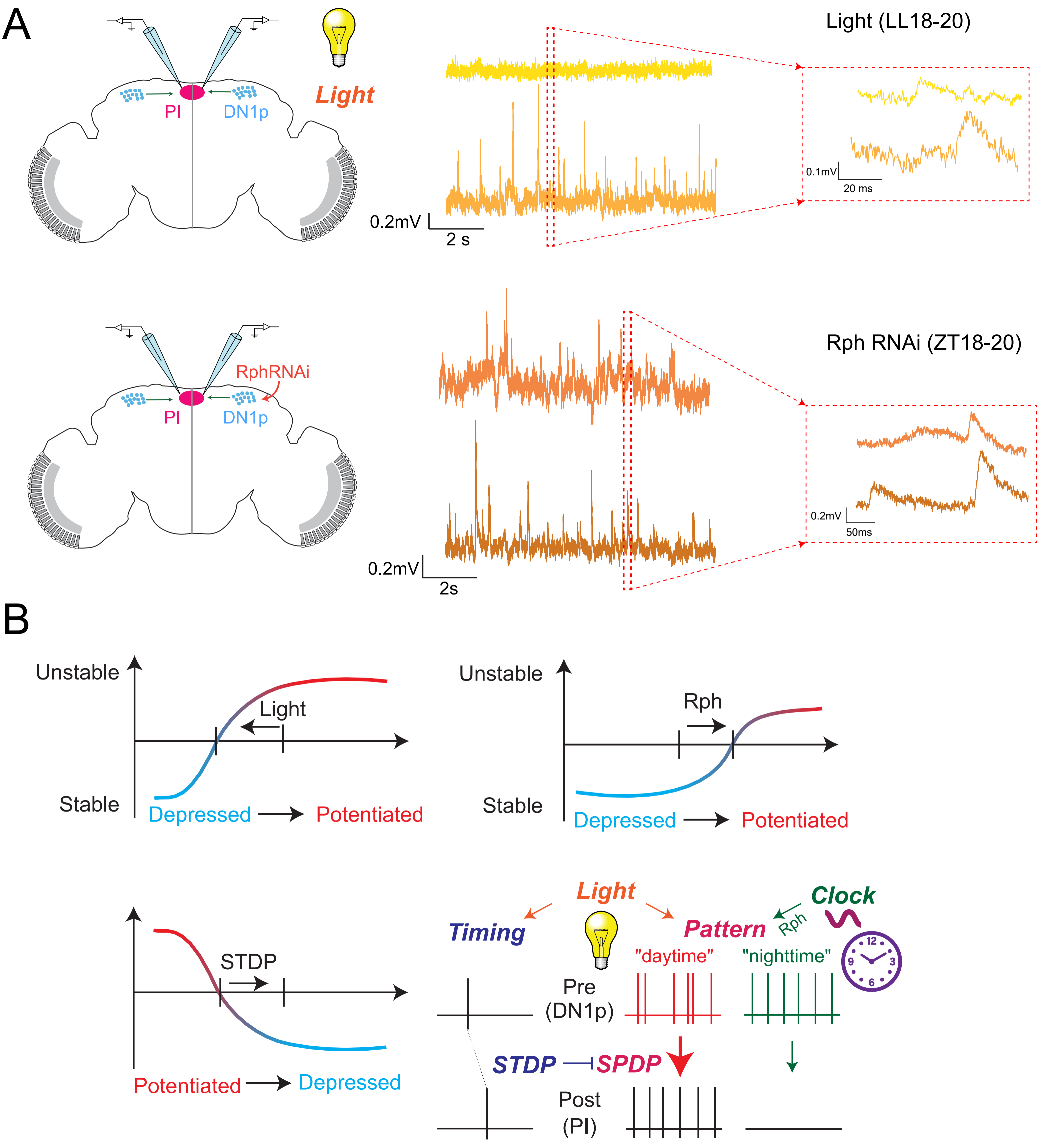
